# Linked origins but distinct roles for extreme length and sequence variation at a tandem repeat in *CACNA1C*

**DOI:** 10.64898/2026.04.23.720446

**Authors:** Janet H.T. Song, Vivien Zhao, Rachel L. Grant, David M. Kingsley

**Affiliations:** Department of Human Evolutionary Biology, Harvard University, Cambridge, MA, 02138, USA; Department of Developmental Biology, Stanford University, Stanford, CA, 94305, USA; Howard Hughes Medical Institute, Stanford University, Stanford, CA, 94305, USA

**Author notes:** Corresponding authors *Email addresses:* (Janet H.T. Song), (David M. Kingsley).

**Keywords:** tandem repeat, VNTR, TRACT, sequence composition, length variation, bipolar disorder, schizophrenia, *CACNA1C*, *CACNA1C-IT3*

## Abstract

Length variation in tandem repeats is a well-established driver of disease risk and is commonly assumed to arise from persistent genomic instability. Here, we characterize TRACT, a 30-bp variable number tandem repeat (VNTR) intronic to the calcium channel gene *CACNA1C*. TRACT exhibits extreme length variation (3-30+ kb) and has been previously linked to risk for bipolar disorder and schizophrenia. By examining multiple human cohorts, we find that TRACT alleles are strikingly bimodal in both length and sequence composition. Short alleles (TRACT^S^, ∼6 kb) and long alleles (TRACT^L^, ∼24 kb) are enriched for distinct 30-bp variants and are found on separate haplotypes that arose prior to the human migration out of Africa. Our data suggest that these ancient alleles expanded via perfect repeat tracts that were disrupted by accumulated mutations to result in relative length stability in extant humans, where there is no evidence for overt germline or somatic instability. Interestingly, neuropsychiatric disease risk is associated with specific 30-bp variants within TRACT^S^ alleles, but not with overall TRACT length or with 30-bp variants enriched in TRACT^L^ alleles. Instead, TRACT^L^ alleles are associated with decreased gene expression in fibroblasts and testis. Together, these findings motivate joint examination of both sequence composition and length variation to fully understand the effects of VNTRs on evolution, trait variation, and disease risk.

## Introduction

Tandem repeats have been implicated in a wide range of diseases, including Huntington disease, spinocerebellar ataxia, and epilepsy, and likely contribute to undetected variation and missing heritability for additional human traits and diseases (Hannan, 2018; Chaisson et al., 2023; Lamkin and Gymrek, 2024). For most of the repeat diseases described to date, the associated repeats are short tandem repeats (STR, repeat unit ≤6 bp) where genomic instability results in repeat expansions or contractions that are associated with disease risk (Chaisson et al., 2023). More recent studies have also implicated length variation with disease risk for variable number tandem repeats (VNTR, repeat unit ≥6 bp), including a 69-bp VNTR where repeat expansion is associated with sporadic amyotrophic lateral sclerosis (Course et al., 2020). These findings have established length variation in tandem repeats, mediated through persistent genomic instability, as a potent source of genetic variation underlying disease risk.

In contrast, we recently identified a VNTR where it appears that sequence variation - but not length variation - is associated with disease risk (Song et al., 2018). This VNTR has a 30-bp repeat unit, is intronic to the calcium channel gene *CACNA1C*, and is flanked by SNPs that have been strongly and consistently associated with bipolar disorder and schizophrenia by genome-wide association studies (GWAS) (Ferreira et al., 2008; Ripke et al., 2014; Song et al., 2018). Strikingly, we and others have found that sequence variation in the 30-bp repeat unit is tightly linked to these GWAS marker SNPs (Song et al., 2018; Moya et al., 2025). Mouse and human organoid studies of this VNTR, called TRACT, demonstrate that TRACT affects the neuronal response to stimulation (Song et al., 2025), suggesting that genetic variation in TRACT is a likely causal contributor to neuropsychiatric disease risk at *CACNA1C*.

In addition to differences in sequence composition, TRACT also exhibits extreme length variation among humans. TRACT ranges from hundreds to thousands of copies of 30-bp units (∼ 3-30+ kb overall), with ∼ 80% of alleles around 6 kb in length (Song et al., 2018). Here, we further investigated how length variation in TRACT arises and whether repeat expansion may have functional consequences. Our analyses suggest that extreme length variation at TRACT resulted from two ancient alleles that differed in sequence composition and arose prior to the migration of humans out of Africa. These distinct sequence compositions resulted in two distinct length distributions in extant humans, likely due to differences in the rate of expansion and mutability of specific 30-bp sequence variants. In striking contrast to other tandem repeats with extreme length variation (Semaka et al., 2013; Handsaker et al., 2025), we found no evidence that extant TRACT alleles exhibit substantial germline or somatic copy number variation. Further, we confirmed that TRACT length variation is not associated with neuropsychiatric phenotypes in modern humans, but is associated with gene expression changes in testis and fibroblasts. These findings suggest that both sequence composition and length variation within VNTRs may have significant contributions to human traits and diseases.

## Materials and Methods

### Southern blot evaluation of TRACT length

DNA or tissue from the brain and a non-ectoderm-derived organ (mostly muscle and liver) were obtained from the NIH Neurobiobank and analyzed as described previously (Song et al., 2018). Briefly, 5-10 µg of human DNA was digested with the restriction enzyme BlpI (New England Biolabs), run on a 0.5% agarose gel, transferred, and hybridized at 60^°^C overnight using a radio-labeled probe made from a DNA template with ten 30-bp repeats and ∼ 500 bp of flanking unique sequence (primer set: 5’-AGGAAAGCACCATTCCCCAG-3’ and 5’-CCATCCCTGAGTTGTGTGCA-3’). The next day, the membrane was washed before exposure and subsequent imaging.

### Analysis of 1000 Genomes Project Data

We analyzed individuals sequenced as part of the 1000 Genomes Project (1000 Genomes Project Consortium, 2015) as previously described (Song et al., 2018). 30-bp variants were assigned numbers based on their prevalence in the 1000 Genomes Project (e.g., variant 1 is the most common repeat sequence variant, variant 2 is the second most common, and so on). A list of 30-bp variants referenced in this manuscript is listed in Table S1. Principal components analysis and k-means clustering were performed on the proportions of each of the 292 30-bp variants identified an average of >0.2 times per individual in the 1000 Genomes Project (denominator is the number of repeats in each individual that matches one of the 292 30-bp variants), matching prior analysis methods (Song et al., 2018). To analyze the proportion of 30-bp variants that have a 1 bp difference from a given 30-bp variant (Fig. 3C, Fig. S4), proportions were calculated such that the denominator is all 30-bp repeat units with ≤5 bp difference from variant 1 (GACCCTGACCTGACTAGTTTACAATCACAC) to capture a larger sample of rare variants. Variant 1, which has a 1 bp sequence difference from both variant 2 and variant 3, was excluded from these analyses. Results are similar when only the 292 most prevalent 30-bp variants are analyzed (Song et al., 2018).

Statistical comparisons were performed with the 2-sample Kolmogorov–Smirnov test, the Wilcoxon rank-sum test, or least-squares linear regression, as indicated in figure legends. Significance was assessed after Bonferroni correction. Phasing was performed with Beagle 5.0 (Browning and Browning, 2007), and phased alleles were visualized with Haplostrips (Marnetto and Huerta-Sánchez, 2017). Variants in linkage disequilibrium with TRACT^L^ were identified using VCFtools (Danecek et al., 2011). The GWAS catalog (v1.0.2) was downloaded from the NHGRI-EBI GWAS catalog (https://www.ebi.ac.uk/gwas/docs/file-downloads) in November 2025. There was one GWAS SNP with *r*^2^ > 0.1 with TRACT^L^, rs11609729 (*r*^2^ = 0.82), which is associated with electrocardiogram morphology (Verweij et al., 2020).

### Analysis of long-read assembled genomes from HGSVC, HPRC, and the Platinum Pedigree

Haplotype-resolved assemblies were downloaded from the HGSVC3 FTP site in July 2025 (https://ftp.1000genomes.ebi.ac.uk/vol1/ftp/data_collections/HGSVC3/working/) (Logsdon et al., 2025), HPRC S3 bucket in November 2025 (Liao et al., 2023), and Platinum Pedigree S3 bucket in May 2025 (Kronenberg et al., 2025). For the four individuals in both HGSVC3 and HPRC, assemblies from HGSVC3 were used. TRACT was extracted from each assembly by searching for the top 3 most common 30-bp variants and obtaining the sequence between the expected flanking sequences. Repeat units were identified as previously described (Song et al., 2018). Variant proportions for each allele were calculated by dividing the number of repeat units for each variant by the total number of repeat units in the allele. Alleles with a variant 3 proportion > 0.2 were considered TRACT^L^ alleles, while those with a variant 3 proportion < 0.2 were considered TRACT^S^ alleles.

To ensure the final set of assemblies contained only unrelated individuals, we removed alleles from children from the three HGSVC3 family trios and non-founder alleles from the Platinum Pedigree. Additionally, we used vcftools –relatedness2 (Danecek et al., 2011) to calculate a relatedness statistic for NA19331 and NA19036, which share an identical TRACT^L^ allele. Both alleles were included in the analyses because the relatedness statistic of 0.027 indicated that they were not first-degree relatives, although results do not change if one of these individuals is excluded. Statistical comparisons between TRACT^S^ and TRACT^L^ alleles were performed using the 2-sample Kolmogorov-Smirnov test to compare the distributions of perfect repeat tract lengths and the Wilcoxon rank-sum test to compare the number and proportion of perfect repeat tracts containing at least five 30-bp units. Haplotype-resolved alleles from HGSVC3 were visualized with Haplostrips (Marnetto and Huerta-Sánchez, 2017).

The twelve TRACT alleles from Platinum Pedigree founder individuals were labeled A-L. Global pairwise sequence alignment was used to determine non-founder genotypes. For each parent-child trio, each of the two child alleles was aligned against each of the four parental alleles using PairwiseAligner from Biopython’s Bio.Align module (Cock et al., 2009) with a match score of 1, a mismatch score of -1, an open gap score of -3, and an extend gap score of -0.5. Alignment scores were calculated using the aligner.score() method. The parental allele with the maximum alignment score was selected as the inherited allele. Each child allele was compared with the inherited parental allele using the .counts() method and a mismatch between the sequences was counted as a *de novo* mutation.

### Analysis of data from the Genotype-Tissue Expression Project (GTEx)

DNA sequencing data from the Genotype-Tissue Expression project (GTEx) were obtained from dbGaP accession number phs000424.v8.p2 in April 2020. We downloaded the reads that mapped to chr12:2255491-2256390 (hg38) for each individual in GTEx. Given that TRACT is mis-annotated in the human reference genome (Song et al., 2018), we first confirmed that reads that contain the 30-mer correctly map to these coordinates. We downloaded all of the reads for 8 individuals and searched for reads containing the 30-mer. In 6 of the 8 individuals, all of the reads containing the 30-mer were correctly mapped. In 2 of the 8 individuals, 1 (0.06%) and 5 (0.11%) reads respectively were incorrectly mapped. This indicates that most reads that contain the 30-mer sequence correctly map to TRACT. Determining the proportion of 30-bp variants and phasing TRACT^L^ was performed as described above.

Normalized gene expression matrices and PEER covariates from the GTEx Portal were downloaded in April 2020. To test for an association between gene expression and TRACT sequence composition, we fit a linear model using the R lm function to the proportion of a given 30-bp variant and *CACNA1C* expression, adjusting for all covariates. We tested all genes 1 Mb upstream and downstream of TRACT. 30-bp variants were only significantly correlated with *CACNA1C* expression in the cerebellum and cerebellar hemisphere (GTEx measures the cerebellum twice, as “cerebellum” and “cerebellar hemisphere”). Significance was determined after Bonferroni correction. The same analysis was also run after adjusting for read depth at TRACT (number of reads that contain a given 30-bp variant / total number of reads) or using the raw number, rather than the proportion, of specific 30-bp variants. These input sets obtained similar results to examining the proportion of a given 30-bp variant. We also analyzed the data either with or without individuals containing TRACT^L^ alleles. In most cases, associations were slightly stronger when individuals with TRACT^L^ alleles were excluded, but our findings were unchanged with or without these individuals. In addition, fine-mapping analysis of this region in the third intron of *CACNA1C* suggests that there are at least two eQTL loci for the cerebellum and cerebellar hemisphere, one centered at the BD and SCZ GWAS locus that is the focus of this study and another at rs886898 (Lonsdale et al., 2013; Hormozdiari et al., 2014; Brown et al., 2017; Wen et al., 2017). The *r*^2^ between rs886898 and the neuropsychiatric GWAS SNPs ranges from 0.0016 to 0.0004, suggesting that these two loci are not linked. Including the genotype at rs886898 as a covariate improved association strength but did not change our findings. The results shown in Fig. 4 and Fig. S8 use the proportion of specific 30-bp variants without adjusting for read coverage, exclude individuals with TRACT^L^, and do not include rs886898 as a covariate.

To test for an association between gene expression and TRACT length, we coded each individual as TRACT^S/S^, TRACT^S/L^, and TRACT^L/L^ based on their SNP haplotype and used FastQTL (Ongen et al., 2016) to identify associated gene expression changes. As an orthogonal approach, we also fit a linear model using the R lm function between gene expression and TRACT read depth (number of reads that map to TRACT / total number of reads). All covariates from GTEx were included, and significance was determined after Bonferroni correction. Findings were highly concordant regardless of method. The results shown in Fig. 4 and Fig. S8 are from FastQTL analysis. Unadjusted p-values are plotted in Fig. 4 and Fig. S8.

## Results

### TRACT alleles are distinguished by both length variation and sequence composition

Extant TRACT alleles vary in length, ranging from 3-30+ kb (100-1000+ repeat units), and in sequence, where different repeat units contain variants on a common 30-bp motif. Strikingly, we found that both length and sequence variation at TRACT is bimodal. In a cohort of 246 individuals analyzed by Southern blot for TRACT length (181 individuals from (Song et al., 2018) and 65 individuals added in this study), 95% of TRACT alleles are centered at ∼ 6 kb in length, while the remaining 5% of alleles are centered at ∼ 24 kb in length (Fig. 1A; Fig. S1A). No TRACT alleles were identified between 15-23 kb in this cohort, emphasizing the bimodality of the TRACT length distribution.

**Figure 1:**
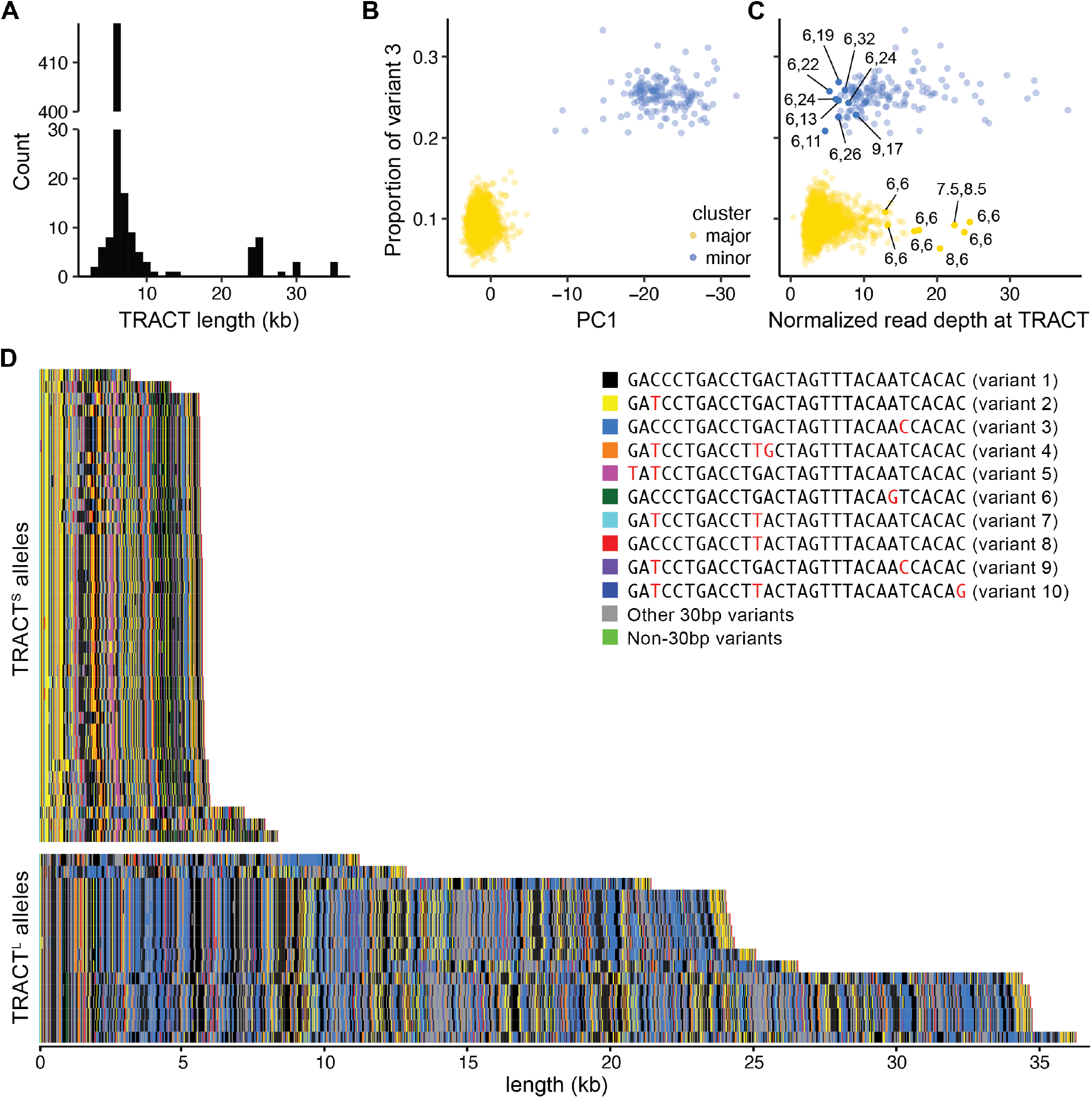
TRACT length variation is linked to sequence variation. (A) Distribution of TRACT length by Southern blot for *N* = 492 alleles (362 alleles from (Song et al., 2018) and 130 alleles added in this study). (B) For individuals in the 1000 Genomes Project (1000 Genomes Project Consortium, 2015), principal component (PC) analysis of 30-bp variant proportions separates individuals by variant 3 proportion along PC1. K-means clustering of 30-bp variant proportions identified a major cluster (yellow; 94% of individuals) and a minor cluster (blue; 6% of individuals). Variant 3 proportion is significantly different between the major and minor clusters (*p* < 10^−96^, Wilcoxon rank-sum test). (C) Individuals in the 1000 Genomes Project with high variant 3 proportion are enriched for increased read depth (*p* < 10^−40^, 2-sample Kolmogorov–Smirnov test). The x-axis is the number of reads that map to TRACT divided by the total number of reads in each sample *x*10^6^. TRACT allele lengths are listed for 17 individuals assessed by Southern blot. (D) Visualization of a representative sample of 40 TRACT^S^ and 16 TRACT^L^ alleles from HGSVC (Logsdon et al., 2025), HPRC (Liao et al., 2023), and PP (Kronenberg et al., 2025). The ten most common 30-bp variants are represented by different colors. TRACT alleles from HGSVC, HPRC, and PP separate by sequence and length into TRACT^S^ alleles (*N* = 574; median length = 5,932 bp; standard deviation = 1,045 bp) and TRACT^L^ alleles (*N* = 18; median length = 27,798 bp; standard deviation = 11,653 bp) that correspond to the major and minor clusters in Fig. 1B, respectively.

TRACT alleles also separate into two clusters based on sequence composition. Although each 30-bp repeat unit in TRACT is based on the same recognizable motif, SNPs within the 30-bp repeat unit are common. For instance, the most common 30-bp unit (variant 1) makes up 31% of all repeat units and has the sequence 5’-GACCCTGACCTGACTAGTTTACAATCACAC-3’ (Song et al., 2018) (Table S1). The second most frequent 30-bp unit (variant 2) makes up 17% of repeat units and has the sequence 5’-GATCCTGACCTGACTAGTTTACAATCACAC-3’ (difference from variant 1 is underlined). To interrogate whether TRACT alleles are distinguished by sequence composition, we determined the proportion of specific 30-bp variants in TRACT in individuals in the 1000 Genomes Project using short-read DNA sequencing data (1000 Genomes Project Consortium, 2015). Principal components analysis (PCA) of the proportions of different 30-bp variants separated individuals into two distinct clusters along PC1 (Fig. 1B; Fig. S1B). These clusters were also recovered by k-means clustering, with 94% of individuals in the major cluster and 6% of individuals in the minor cluster.

Of the 30-bp units with the strongest contribution to PC1, variant 3 (GACCCTGACCTGACTAGTTTACAACCACAC, difference from variant 1 is underlined) cleanly separated the major and minor clusters (Fig. 1B). Variant 3 makes up 9.6% ±1.4% of repeat units in individuals in the major cluster but 25.3% ±2.0% of repeat units in individuals in the minor cluster (*p* < 10^−96^ by the Wilcoxon rank-sum test) (Table S2). We used low (∼ 10%) and high (∼ 25%) variant 3 proportion as a proxy for the major and minor clusters, respectively.

The striking bimodality of both length and sequence variants in TRACT – and the similar proportions of major and minor clusters – raised the hypothesis that length and sequence variation in TRACT are linked. To test this hypothesis, we examined normalized read depth in individuals from the 1000 Genomes Project as a proxy for TRACT length (Materials and Methods). Although high variant 3 proportion is significantly correlated with increased read depth in the 1000 Genomes Project (*p* < 10^−40^ by 2-sample Kolmogorov-Smirnov test), there is substantial overlap of read depth distributions between individuals with low and high variant 3 proportions (Fig. 1C). To investigate how tightly length is associated with sequence composition, we performed Southern blot for TRACT in 17 individuals that would most strongly contradict this association (filled circles in Fig. 1C): eight individuals with high read depth but low variant 3 proportion, and nine individuals with low read depth but high variant 3 proportion. Strikingly, we found that all tested individuals with low variant 3 proportion had two alleles of ∼6 kb in size, and all tested individuals with high variant 3 proportion had one allele that is 11+ kb. This suggests that overlapping read depth distributions between high and low variant 3 proportion clusters is likely due to noise in short-read DNA sequencing and that the bimodal distribution of TRACT length observed by Southern blot likely directly corresponds to distinct sequence compositions.

To gain allele-level resolution of TRACT, we next examined the Human Genome Structural Variation Consortium (HGSVC) (Logsdon et al., 2025), the Human Pangenome Reference Consortium (HPRC) (Liao et al., 2023), and the Platinum Pedigree (PP) (Kronenberg et al., 2025), where fully assembled genomes from long-read DNA sequencing are available for 296 unrelated individuals. Visualization of TRACT alleles by coloring the ten most common variants (Fig. 1D; Fig. S1C) confirmed that TRACT alleles contain two distinct sequence compositions that correspond to differences in TRACT length. 96.96% of TRACT alleles (*N* = 574) have a sequence composition that is characterized by a high proportion of variant 2 (yellow in Fig. 1D; 18.69% ± 2.42%) and a low proportion of variant 3 (blue in Fig. 1D; 10.76% ± 1.07%). These alleles correspond to the shorter length distribution identified by Southern blot (median length = 5,932 bp; standard deviation = 1,045 bp). The remaining 3.04% of TRACT alleles (*N* = 18) have a distinct sequence composition, which is characterized by a low proportion of variant 2 (7.26% ± 0.84%) and a high proportion of variant 3 (28.94% ± 1.01%). These alleles correspond to the longer length distribution identified by Southern blot (median length = 27,798 bp; standard deviation = 11,653 bp). We refer to these distinct sets of alleles as TRACT^S^ and TRACT^L^, respectively. Combined with our findings in the 1000 Genomes Project, these results demonstrate that TRACT alleles separate into two clusters with distinct sequence compositions and length distributions.

### TRACT^L^ arose once in the human lineage

Given that TRACT^S^ and TRACT^L^ are distinct in sequence and length, we wondered whether TRACT^L^ alleles arose on a single haplotype in the human lineage or are the result of recurrent repeat expansions on different haplotypes. We coded TRACT length as a variant in individuals from the 1000 Genomes Project and phased the 2 Mb interval surrounding TRACT (Materials and Methods). Individuals with low variant 3 proportion are TRACT^S/S^. We coded all individuals with high variant 3 proportion as heterozygous (TRACT^S/L^) because only 4-5 individuals are expected to be homozygous (TRACT^L/L^) in the 1000 Genomes Project under Hardy-Weinberg equilibrium (Song et al., 2018).

TRACT^L^ alleles cleanly clustered together based on flanking sequence variants, suggesting that TRACT^L^ arose on only one haplotype (Fig. S2). Consistent with our estimate that 4-5 individuals would be homozygous for TRACT^L^ under Hardy-Weinberg equilibrium, both alleles from five individuals with high variant 3 proportion clustered with TRACT^L^ alleles (red arrowheads in Fig. S2). When these five individuals were recoded as homozygous for TRACT^L^, 135 SNPs had *r*^2^ > 0.5 with TRACT^L^, of which 51 SNPs had *r*^2^ > 0.9 and 5 SNPs were in complete linkage disequilibrium (*r*^2^ = 1) (Table S3). We confirmed that TRACT^L^ alleles are found on only one haplotype by examining HGSVC, which is a smaller cohort (*N* = 65) but provides allelic resolution due to long-read sequencing. TRACT^L^ alleles in HGSVC clustered together and were found on the same haplotype identified in individuals from the 1000 Genomes Project (Fig. 2).

**Figure 2:**
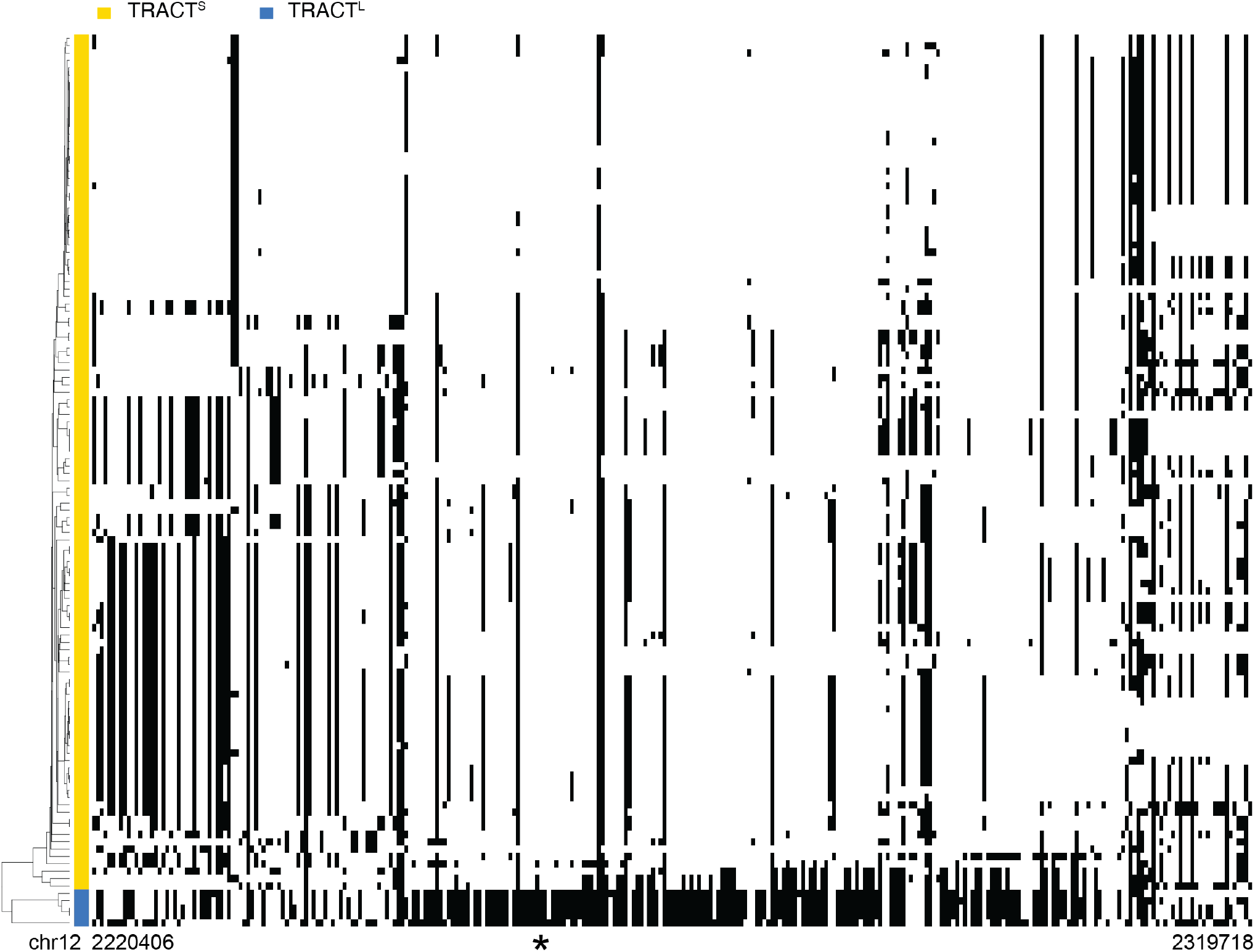
TRACT^L^ is found on one haplotype in extant humans. SNPs in haplotype-resolved alleles from HGSVC were clustered for the interval surrounding TRACT (chr12: 2220406-2319718, hg38). The genomic position of TRACT is indicated by an asterisk. TRACT^S^ (yellow) and TRACT^L^ (blue) alleles clustered separately.

To date the emergence of TRACT^L^, we examined which populations in the 1000 Genomes Project contained TRACT^L^ alleles. TRACT^L^ is found in all super populations: Africans, Ad-Mixed Americans, East Asians, Europeans, and South Asians (Fig. S3). Together, these results suggest that TRACT^L^ is derived from a single ancestral allele that arose prior to the human migration out of Africa.

### Stretches of perfect repeat tracts likely drove TRACT expansion

Tandem repeat instability has been strongly associated with the length of perfect repeat tracts, defined as regions containing only repeat units of the same sequence (Eichler et al., 1994; Kunst and Warren, 1994; Cumming et al., 2018; Lee et al., 2019). We therefore hypothesized that differences in length between TRACT^S^ and TRACT^L^ alleles may be derived from differences in the lengths of perfect repeat tracts. Indeed, we found that TRACT^L^ alleles are enriched for perfect repeat tracts compared to TRACT^S^ alleles (*p* < 10^−49^ by the 2-sample Kolmogorov-Smirnov test, Fig. 3A), with both a larger number of perfect repeat tracts (*p* < 10^−20^ by the Wilcoxon rank-sum test) and a higher proportion of alleles comprised of perfect repeat tracts (*p* < 10^−11^ by the Wilcoxon rank-sum test) (Fig. 3B). Most perfect repeat tracts that were composed of at least five repeat units were of variants 1, 2, or 3. Of note, the shortest TRACT^L^ alleles (11.2 kb; asterisk in Fig. 3A) had much longer perfect repeat tracts than the other TRACT^L^ alleles; their longest perfect repeat tracts comprised 26 units of variant 3. Given that TRACT^L^ alleles <24 kb are rarely observed (Fig. 1A), this suggests that short TRACT^L^ alleles are enriched for long perfect repeat tracts that are prone to repeat expansion, resulting in their low frequency in extant humans. Further, this suggests that the observed TRACT^S^ and TRACT^L^ length distributions in extant humans likely arose from a balance between the rate of expansion of perfect repeat tracts, mutations that arise to disrupt them, and differences in these rates for specific 30-bp variants.

**Figure 3:**
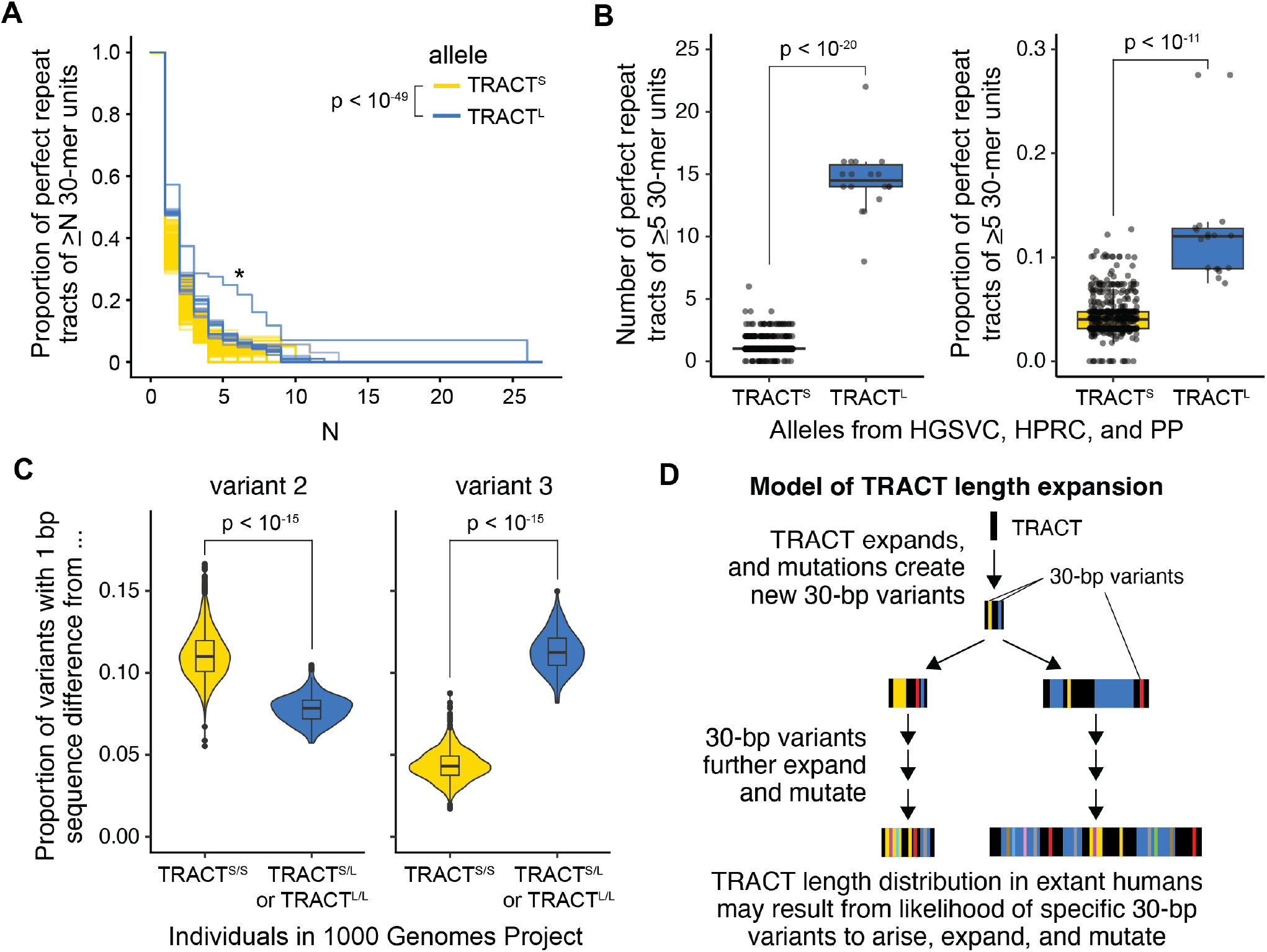
TRACT^L^ is characterized by long perfect repeat tracts and increased variant 3 proportion. (A) TRACT^L^ alleles (blue; *N* = 18) are enriched for longer perfect repeat tracts, defined as contiguous stretches of the same 30-bp variant, compared to TRACT^S^ alleles (yellow; *N* = 574) for alleles from HGSVC, HPRC, and PP (*p* < 10^−49^, 2-sample Kolmogorov-Smirnov test). (B) Number (left; *p* < 10^−20^, Wilcoxon rank-sum test) and proportion (right; *p* < 10^−11^, Wilcoxon rank-sum test) of perfect repeat tracts of at least 5 30-mer units differs between TRACT^S^ and TRACT^L^ alleles. (C) In the 1000 Genomes Project, TRACT^S/S^ individuals (*N* = 2334) are enriched for 30-mer variants with a 1 bp sequence difference from variant 2 (*p* < 10^−15^, Wilcoxon rank-sum test) and depleted for 30-mer variants with a 1 bp sequence difference from variant 3 (*p* < 10^−15^, Wilcoxon rank-sum test) when compared to TRACT^S/L^ and TRACT^L/L^ individuals (*N* = 156). Variant 1, which is the most prevalent 30-bp variant and has a 1 bp sequence difference from both variant 2 and variant 3, was excluded from these analyses. (D) Model for how TRACT length distribution in extant humans may have arisen.

To determine how differences in the propensity of distinct 30-bp variants to expand or mutate may have contributed to the observed length and sequence distributions at TRACT, we further investigated how the distinct sequence compositions of TRACT^S^ and TRACT^L^ alleles may have arisen. The most common 30-bp variant in all TRACT alleles is variant 1 (GACCCTGACCTGACTAGTTTACAATCACAC, black in Fig. 1D). Variant 1 has a 1 bp difference from variant 2 (GATCCTGACCTGACTAGTTTACAATCACAC, yellow in Fig. 1D), the next most common variant in TRACT^S^ alleles. Variant 1 also has a 1 bp difference from variant 3 (GACCCTGACCTGACTAGTTTACAACCACAC, blue in Fig. 1D), the next most common variant in TRACT^L^ alleles. Given that new variants are likely generated by mutating existing variants, this suggests that variants 2 and 3 arose from 1 bp mutations in variant 1, and that the establishment of either variant 2 or 3 as a high frequency variant may have driven subsequent differences in sequence and length composition between TRACT^S^ and TRACT^L^ alleles.

If extant TRACT^S^ and TRACT^L^ sequence diversity originates from variants 2 and 3 respectively, 30-bp variants with a 1 bp sequence difference from variant 2 will be enriched in individuals with TRACT^S^ alleles, and 30-bp variants with a 1 bp sequence difference from variant 3 will be enriched in individuals with TRACT^L^ alleles. Consistent with this prediction, we found that individuals with a TRACT^L^ allele (TRACT^S/L^ or TRACT^L/L^) in the 1000 Genomes Project are enriched for variants that are 1 bp from variant 3 and that this enrichment holds when only considering rare variants (<1%, <0.1%, or <0.01% frequency) (Fig. 3C, Fig. S4). Similarly, variants that are 1 bp from variant 2 are enriched in TRACT^S/S^ individuals (Fig. 3C, Fig. S4). These results suggest that divergent sequence compositions between TRACT^S^ and TRACT^L^ alleles originated from variants 2 and 3 respectively.

Together, these observations suggest a model whereby TRACT expanded via perfect repeat tracts (Fig. 3D), similar to other previously described disease-associated tandem repeats (Eichler et al., 1994; Kunst and Warren, 1994; Cumming et al., 2018; Lee et al., 2019). As perfect repeat tracts expanded over time, mutations arose, introducing new 30-bp variants that decreased the length of perfect repeat tracts and slowed further TRACT expansion. Given the strong modality of extant TRACT^S^ and TRACT^L^ alleles at lengths of ∼6 kb and ∼24 kb respectively, relative stability at these high frequency lengths could have arisen due to differences in starting lengths of perfect repeat tracts and in the propensity of specific 30-bp variants to arise, expand, and mutate.

### No evidence for elevated germline or somatic mutation rate at TRACT

Germline and somatic genomic instability via *de novo* repeat expansions or contractions is a hallmark of disease-associated tandem repeats (Khristich and Mirkin, 2020; Handsaker et al., 2025). To examine whether extant TRACT alleles exhibit germline instability, we examined the Platinum Pedigree (PP), a 4-generation pedigree of 23 individuals with long-read assembled genomes (Porubsky et al., 2025) (Fig. S5). Of the 8 TRACT alleles present in the first generation (G1), seven are TRACT^S^ alleles (lettered A, B, D-H in Fig. S5: *∼*6 kb) and one is a TRACT^L^ allele (lettered C in Fig. S5: 24.2 kb). The 4 additional TRACT alleles (lettered I-L in Fig. S5) introduced through marriage in the third generation are all TRACT^S^ alleles (*∼*6 kb). Strikingly, there are no *de novo* length or sequence changes in TRACT in this pedigree, suggesting that germline mutations in TRACT are not common.

To examine whether TRACT exhibits somatic mosaicism in repeat length, we examined DNA from matched brain and non-ectoderm tissue for 112 individuals, including 50 individuals with schizophrenia (Materials and Methods). TRACT lengths in these individuals ranged from 4-30 kb (Fig. S6). We did not observe any evidence at the resolution of Southern blot for somatic length mosaicism in any individual. Additionally, although TRACT is highly unstable in bacteria (Song et al., 2018), we found that TRACT length in different human cell lines, including HEK293T and multiple induced pluripotent stem cell lines, are within the range observed in modern humans and are stable across passages. This suggests that TRACT is more somatically stable in humans than other disease-associated tandem repeats described to date, which experience levels of somatic length mosaicism readily detectable by Southern blot (Martorell et al., 1998; Beck et al., 2013). Together, these data further support our proposed model where a balance between repeat expansion and mutation accumulation from two starting sequence compositions resulted in relatively stable bimodal distributions centered at *∼*6 kb and *∼*24 kb in extant humans (Fig. 3D).

### TRACT length variation is associated with gene expression changes in fibroblasts and testis

A combination of computational and experimental approaches has established TRACT sequence variation as a likely causative variant at a top GWAS locus for bipolar disorder and schizophrenia (Song et al., 2018; Moya et al., 2025; Song et al., 2025). Intriguingly, variant 3, which is enriched in TRACT^L^ alleles, is one of the 30-bp variants that has been previously linked to the nearby GWAS SNPs (Song et al., 2018). Although this raises the possibility that length variation may also contribute to neuropsychiatric disease risk at this locus, TRACT length has not been previously associated with GWAS SNPs at this locus (Song et al., 2018; Moya et al., 2025). Additionally, we found that the haplotype containing TRACT^L^ (Fig. 2; Fig. S2) is not in linkage disequilibrium with known GWAS SNPs associated with risk for bipolar disorder or schizophrenia (*r*^2^ < 0.1). This makes it highly unlikely that TRACT^L^ – or variant 3, which is much more prevalent in TRACT^L^ than TRACT^S^ (Fig. 1B) – affects neuropsychiatric disease risk.

The previously reported difference in variant 3 proportion between individuals with risk GWAS SNPs (10.7% ± 1.5%) and individuals with protective GWAS SNPs (9.3% ± 1.6%) (Song et al., 2018) is well within the range of variant 3 proportions in the 96% of individuals that are homozygous for TRACT^S^ alleles (Fig. 1B). This suggests that variant 3 proportion may merely be linked to the causative variants underlying disease risk in TRACT^S^ alleles. There are nine other 30-bp variants previously linked to these GWAS SNPs: five (variants 2, 10, 11, 15, 32) are associated with the protective GWAS haplotype and four (variants 13, 27, 33, 44) are associated with the risk GWAS haplotype (Song et al., 2018).

To further disentangle the causality of individual 30-bp variants, we examined their effects on *CACNA1C* expression. GWAS SNPs at this locus are expression quantitative trait loci (eQTL) for *CACNA1C* expression in the cerebellum across multiple cohorts (Gershon et al., 2014; Roussos et al., 2014; Lu et al., 2023; Moya et al., 2025), including the Genotype-Tissue Expression Project (GTEx) (Lonsdale et al., 2013) (Fig. S7). When we examined the association between the proportion of specific 30-bp variants and *CACNA1C* expression in the cerebellum in GTEx, variant 3 proportion was not a significant eQTL (Fig. 4A; Fig. S8). In contrast, other risk- and protective-associated 30-bp variants (2, 10, 13, 27, 33, 44) are significant eQTLs. This suggests that these other 30-bp variants likely drive the contribution of this locus to neuropsychiatric disease risk and that variant 3 proportion is simply a linked variant within TRACT^S^ alleles.

**Figure 4:**
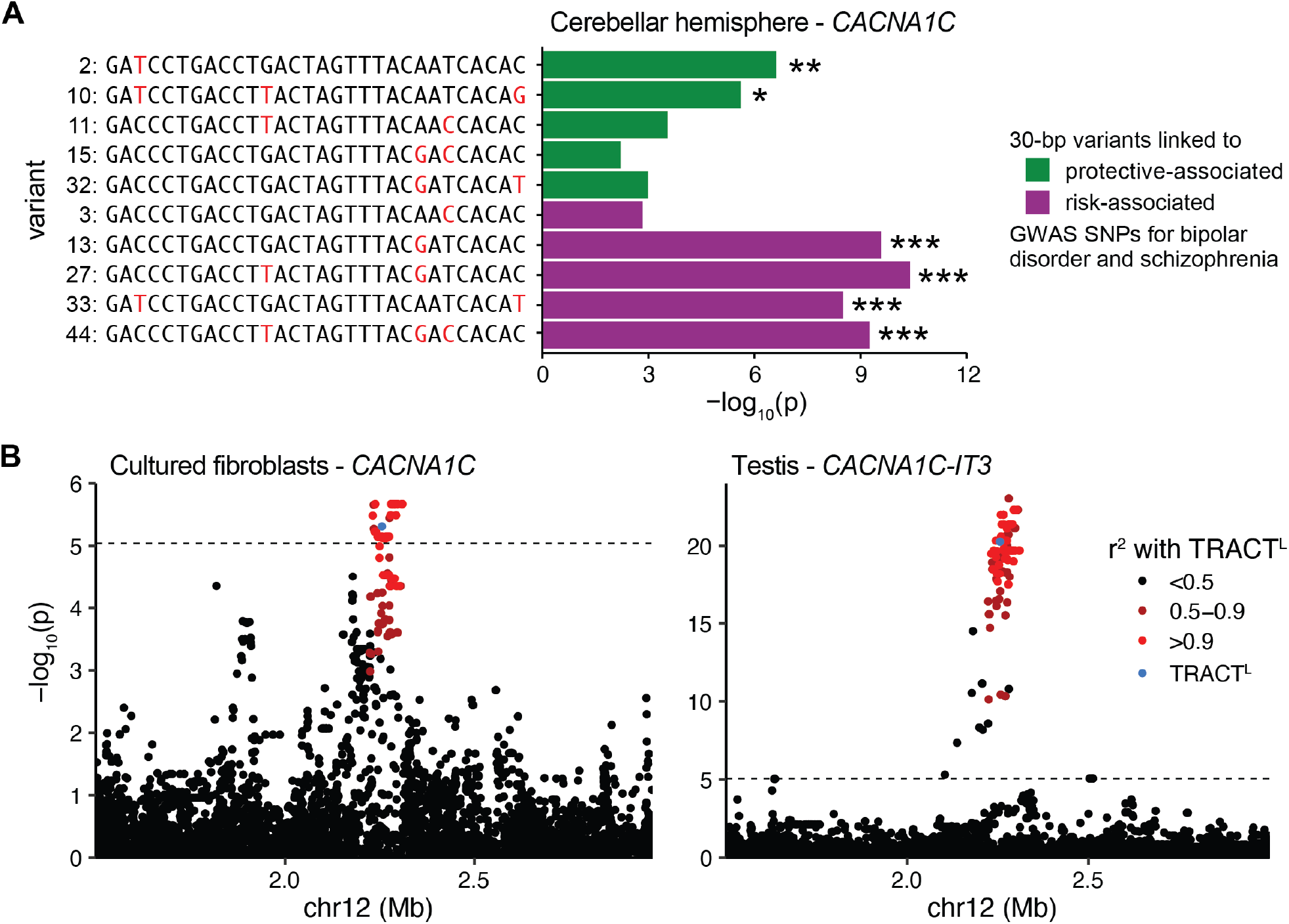
TRACT^L^ is associated with gene expression changes in fibroblasts and testis. (A) In GTEx (Lonsdale et al., 2013), the proportion of variants 2, 10, 13, 27, 33, and 44 are eQTLs for *CACNA1C* expression in the cerebellar hemisphere. *: adjusted *p* < 0.05; **: adjusted *p* < 0.01; ***: adjusted *p* < 10^−4^. (B) TRACT^L^ (blue) and SNPs in linkage disequilibrium with TRACT^L^ (red) are eQTLs for *CACNA1C* expression in cultured fibroblasts (left) and for *CACNA1C-IT3* expression in the testis (right). The dotted line indicates the significance threshold after Bonferroni correction.

What then, if anything, are the functional consequences of TRACT length variation within extant humans? TRACT^L^ is not linked (*r*^2^ > 0.9) to any SNPs in the NHGRI-EBI GWAS catalog (Cerezo et al., 2025), indicating that TRACT^L^ is not associated with any disease risk loci identified to date (Materials and Methods). To determine whether TRACT^L^ may affect nearby gene expression, we examined whether TRACT^L^ is associated with expression changes in genes within 1 Mb of TRACT for each of the 49 tissues assayed in GTEx (Lonsdale et al., 2013). Consistent with the 1000 Genomes Project and HGSVC, TRACT^L^ alleles in GTEx have a distinct sequence composition and are located on one haplotype, allowing us to accurately code TRACT^S^ and TRACT^L^ as a variant in GTEx (Fig. S7).

We found that TRACT^L^ is associated with decreased expression of *CACNA1C* in cultured fibroblasts and with decreased expression of *CACNA1C-IT3*, a long non-coding RNA located 15 kb downstream from TRACT, in the testis (Fig. 4B). TRACT^L^ is not associated with changes in gene expression in the other 47 tissues including the 12 assayed brain regions. Similar results were observed for SNPs in strong linkage disequilibrium with TRACT^L^ (Fig. 4B) or when correlating gene expression changes with TRACT read depth as a proxy for repeat length (Materials and Methods). Together, these results suggest that TRACT length does not have overt effects on neural expression within modern humans, but may mediate expression changes in *CACNA1C* and *CACNA1C-IT3* in fibroblasts and the testis, respectively.

## Discussion

Our analyses suggest that extreme TRACT length variation in modern humans largely arose from ancient sequence divergence rather than from major ongoing genomic instability. Distinct TRACT^S^ and TRACT^L^ alleles arose prior to the human migration out of Africa and differ in both the length of perfect repeat tracts and their 30-bp variant composition. Combined with the lack of overt genomic or somatic instability at TRACT, these findings suggest that TRACT^L^ alleles arose once in the human lineage and that the observed length variation at TRACT resulted from two ancient sequence compositions that differed in rates of expansion and mutation and ultimately stabilized at two disparate length distributions in extant humans.

Our data suggest that ancient expansion of TRACT was driven by stretches of perfect repeat tracts, consistent with established mechanisms of tandem repeat instability (Cumming et al., 2018; Eichler et al., 1994; Kunst and Warren, 1994; Lee et al., 2019). TRACT^L^ alleles are significantly enriched for longer perfect repeat tracts compared to TRACT^S^ alleles, likely reflecting the mechanism that drove expansion to ∼24 kb. Intriguingly, the shortest TRACT^L^ alleles we observed (∼11 kb) contain substantially longer stretches of perfect repeat tracts than longer TRACT^L^ alleles. Because TRACT^L^ alleles <24 kb range are rarely observed (Fig. 1A), we hypothesize that TRACT^L^ alleles in this smaller size range may be particularly prone to expansion due to having longer perfect repeat tracts, and that the observed ∼24 kb modal length represents a length at which mutation accumulation has sufficiently disrupted perfect repeat tracts to reduce or halt further TRACT expansion. The enrichment of variants that are likely derived from variant 2 for TRACT^S^ or from variant 3 for TRACT^L^ further supports this model that sequence-specific differences in the rates of expansion and mutation likely resulted in the observed length distributions in extant humans.

The molecular mechanisms underlying TRACT expansion remain to be elucidated. Potential causes include the formation of secondary DNA structures, replication slippage, and the formation of RNA:DNA hybrids during transcription, all of which have been shown to contribute to repeat instability in STRs (Khristich and Mirkin, 2020). A recent study also suggests that tandem duplication of long stretches of 30-bp units via recombination events may have contributed to TRACT expansion (Moya et al., 2025). Experiments examining the conditions under which TRACT expands, and why TRACT alleles enriched for variant 2 or variant 3 have differing propensities to expand, may provide new insights into general mechanisms of repeat expansion.

For most previously characterized tandem repeat diseases, length variation is an ongoing process of germline and somatic expansion (Khristich and Mirkin, 2020). In contrast, we find that TRACT^L^ alleles appear on only one haplotype across multiple cohorts. This haplotype predates the migration of modern humans out of Africa. In addition, we do not observe germline or somatic length mosaicism at TRACT. We note, however, that the Platinum Pedigree has a small sample size (*N* = 23) and Southern blots are limited in resolution. Although our analyses suggest that there is no overt genomic instability at TRACT comparable to previously described disease-associated repeats (Khristich and Mirkin, 2020; Handsaker et al., 2025), future analyses of additional pedigrees and samples will be needed to determine the exact level of genomic stability at TRACT in extant humans.

Further, we found that the TRACT^L^ haplotype is linked to expression changes in *CACNA1C-IT3* in the testis and *CACNA1C* in cultured fibroblasts. *CACNA1C-IT3* is a functionally uncharacterized human-specific lncRNA whose expression is restricted to the testis in adult humans (Lonsdale et al., 2013). Similarly, although *CACNA1C* function has been extensively studied in the brain and in the heart (Harvey and Hell, 2013; Hofmann et al., 2014; Nanou and Catterall, 2018), little is known about its function in fibroblasts. Additional experimen-tation is required to determine the function of *CACNA1C-IT3* in the testis and *CACNA1C* in fibroblasts and how perturbations to their expression might affect downstream phenotypes.

Sequence variation in TRACT has been linked to risk for bipolar disorder and schizophrenia (Song et al., 2018; Lu et al., 2023; Moya et al., 2025). In contrast, we do not find any association between extreme TRACT length expansion and disease risk. This contrasts with canonical repeat expansion disorders where length variation is the primary pathogenic mechanism. To date, investigations on the effect of tandem repeats on disease risk or gene expression have focused on length variation and largely ignored sequence variation, which can be harder to identify and study (Fotsing et al., 2019; Trost et al., 2020; Bakhtiari et al., 2021; Mitra et al., 2021; Mukamel et al., 2021, 2023). However, recent studies suggest that sequence variants in tandem repeats, particularly VNTRs, are associated with gene expression changes (Lu et al., 2023; Moya et al., 2025). This underscores the importance of comprehensively characterizing sequence variation within tandem repeats and avoiding the common assumption that length variation is the primary driver of functional consequences. Further joint examination of sequence and length variation at tandem repeats will be needed to fully understand the role of tandem repeats in evolution, trait variation, and disease risk.

## Acknowledgements

Research reported in this publication was supported in part by National Science Foundation Graduate Research Fellowships and Stanford Graduate Fellowships (JHTS, RLG), a Stanford CEHG Fellowship (JHTS), and NIH/NIMH grant R00MH136290 (JHTS). DMK is an Investigator of the Howard Hughes Medical Institute. The Genotype-Tissue Expression (GTEx) Project was supported by the Common Fund of the Office of the Director of the National Institutes of Health, and by NCI, NHGRI, NHLBI, NIDA, NIMH, and NINDS. The data used for the analyses described in this manuscript were obtained from the GTEx Portal and dbGaP accession number phs000424.v8.p2 in April 2020. DNA or tissue samples were obtained through the NIH Neurobiobank from the University of Maryland Brain and Tissue Bank and the Mt. Sinai Brain Bank. The content is solely the responsibility of the authors and does not necessarily represent the official views of the National Institutes of Health.

## Author Contributions

**Janet H.T. Song**: Conceptualization, Formal analysis, Investigation, Methodology, Resources, Supervision, Visualization, Writing - original draft, Writing - review & editing. **Vivien Zhao**: Formal analysis, Investigation, Methodology, Visualization, Writing - original draft, Writing - review & editing. **Rachel L. Grant**: Investigation. **David M. Kingsley**: Conceptualization, Resources, Supervision, Writing - original draft, Writing - review & editing.

## Data and code availability

Multiple publicly available datasets were analyzed in this work and can be accessed as described in Materials and Methods. This study did not generate novel code. Data generated in this study are available in the supplement.

