## Supplemental Materials for "Linked origins but distinct roles for extreme length and sequence variation at a tandem repeat in *CACNA1C*"

### Supplemental Tables

Table S1: **30-bp variants**
Table S2: **Proportion of 30-bp variants in individuals in 1000 Genomes Project**
Table S3: **SNPs in linkage disequilibrium with TRACT$^{L}$**

### Supplemental Figures

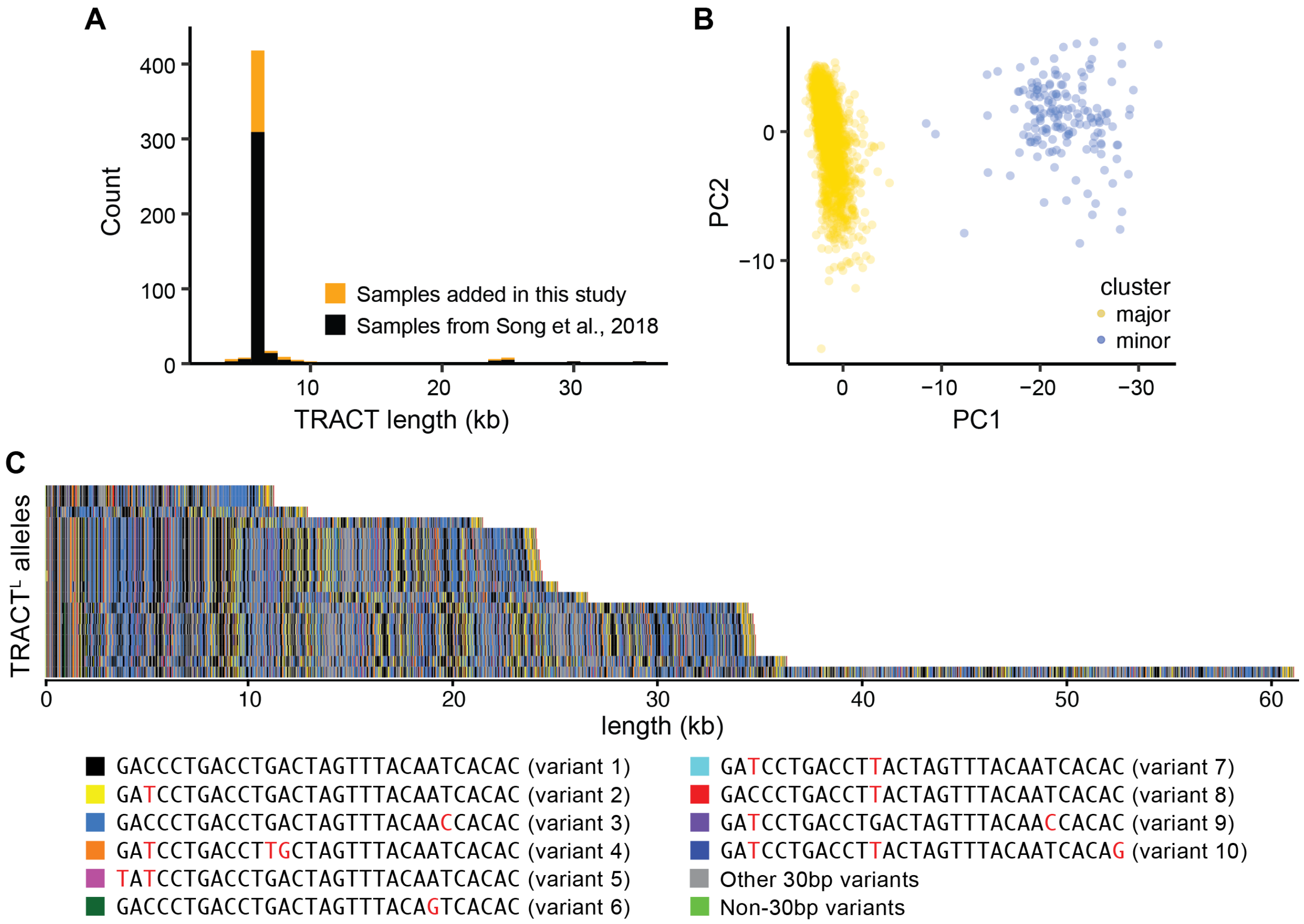


**Figure S1: TRACT length and sequence variation.** (A) Distribution of TRACT length by Southern blot for 362 alleles from (Song et al., 2018) in black and 130 alleles added in this study in orange. (B) PC1 (11.5% of the variance) and PC2 (2.8% of the variance) of 30-bp variant proportions for individuals in the 1000 Genomes Project (1000 Genomes Project Consortium, 2015). (C) Visualization of all TRACT$^L$ alleles from HGSVC (Logsdon et al., 2025), HPRC (Liao et al., 2023), and the Platinum Pedigree (Kronenberg et al., 2025) with coloring of the ten most common 30-bp variants.

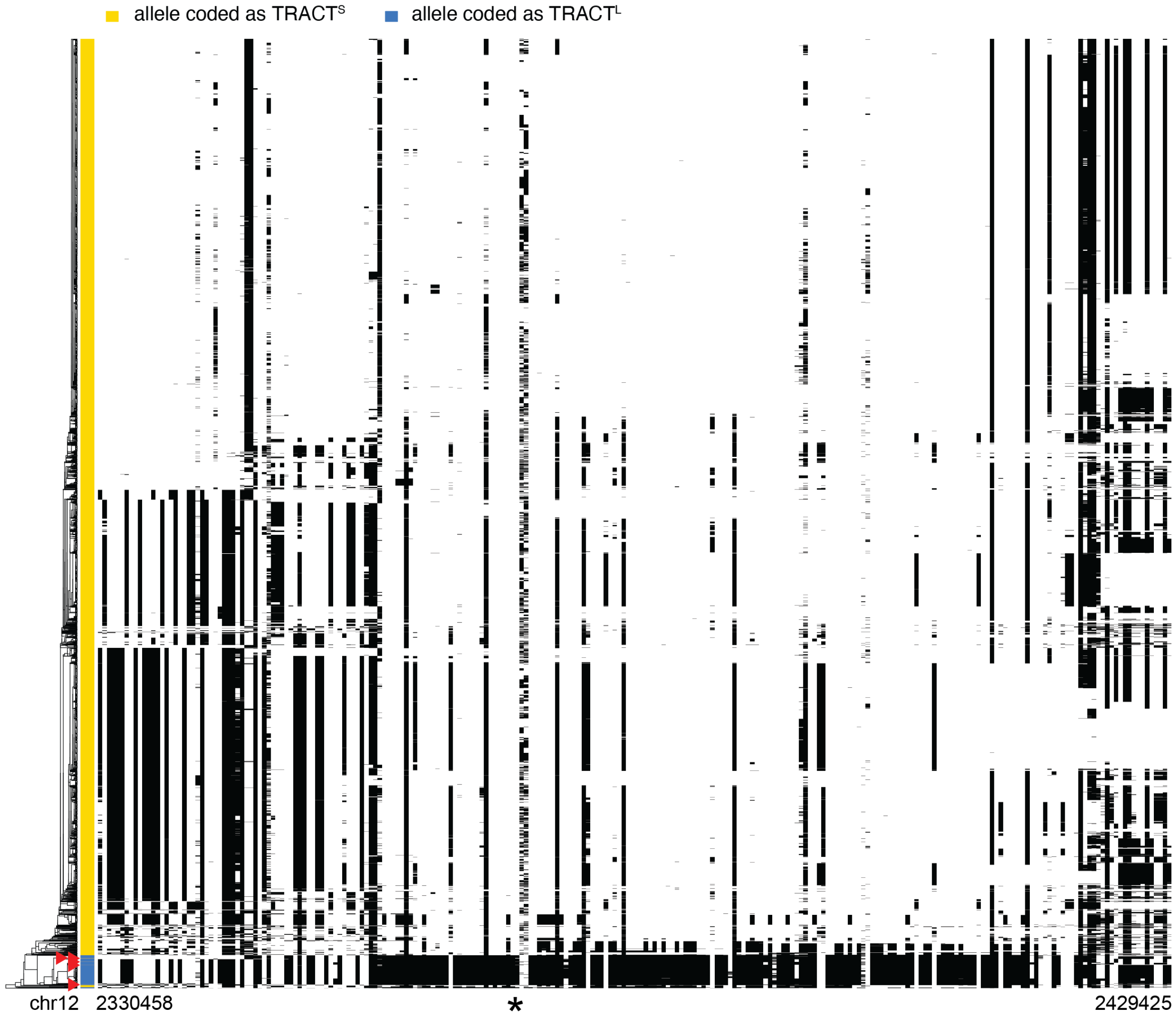


**Figure S2: TRACT$^L$ is found on one haplotype in the 1000 Genomes Project.** TRACT$^S$/TRACT$^L$ was coded as a variant for individuals in the 1000 Genomes Project, and the 2 Mb interval surrounding TRACT was phased using Beagle 5.0 (Browning and Browning, 2007) (Materials and Methods). The phased alleles were visualized using Haplostrips (Marnetto and Huerta-Sánchez, 2017) for chr12:2330458-2429425 (hg19). The genomic position of TRACT is indicated by an asterisk. Five red arrowheads indicate alleles from individuals that are likely homozygous for TRACT$^L$.

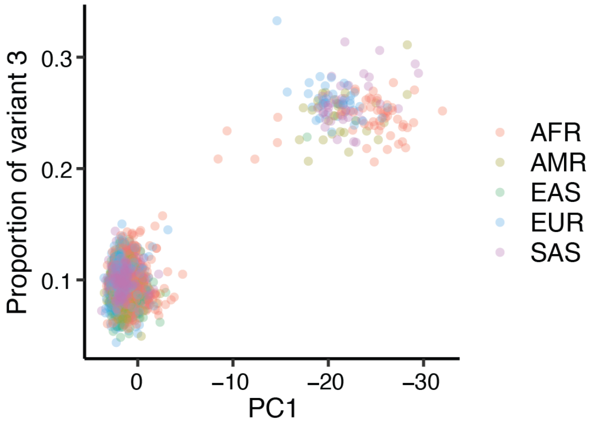


**Figure S3: TRACT$^{L}$ is found in every super population in the 1000 Genomes Project.** Same plot as Fig. 1B colored by super population. AFR: Africans, AMR: Ad-Mixed Americans, EAS: East Asians, EUR: Europeans, SAS: South Asians.

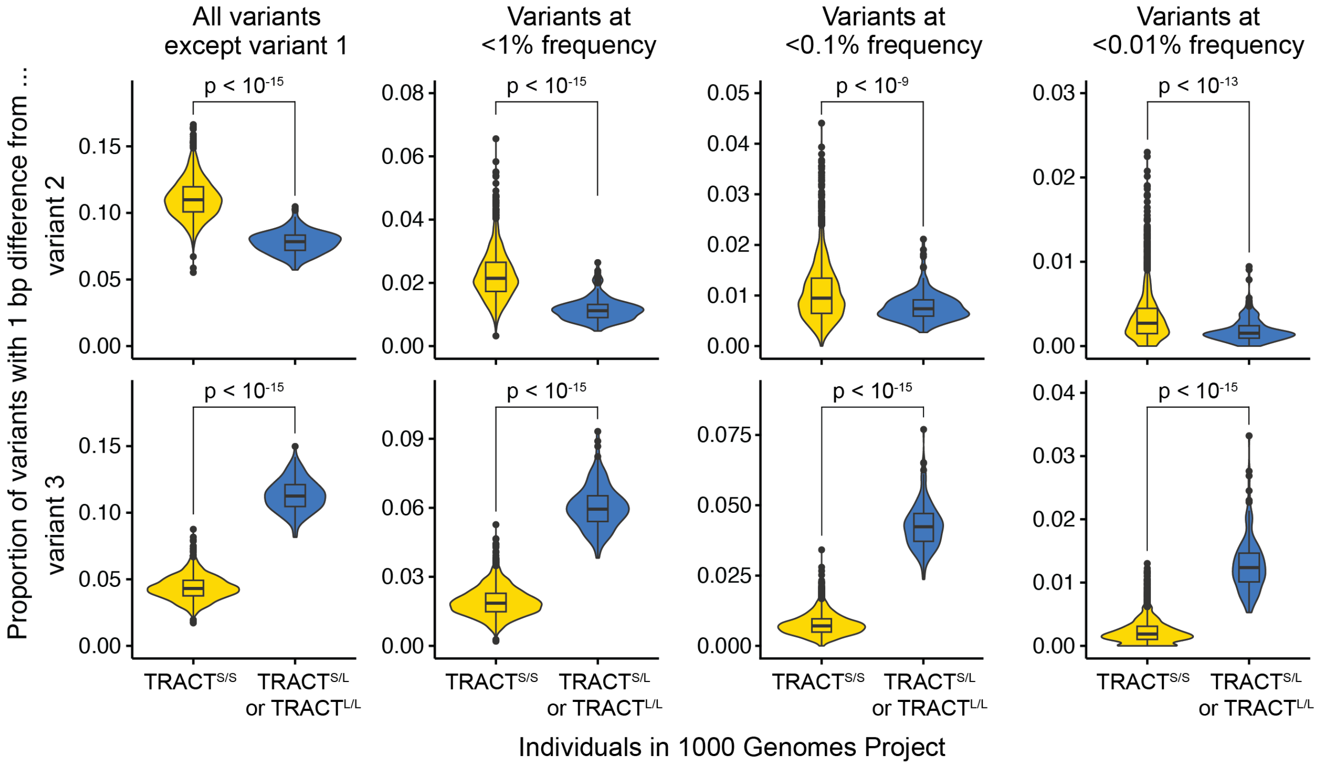


**Figure S4: Proportion of 30-bp variants with 1 bp sequence difference from variant 2 or 3.** Proportion of all variants except variant 1, variants with <1% frequency, variants with <0.1% frequency, and variants with <0.01% frequency that have a 1 bp sequence difference from variant 2 (top) or variant 3 (bottom) for $TRACT^{S/S}$ individuals ($N = 2334$) and $TRACT^{S/L}$ or $TRACT^{L/L}$ individuals ($N = 156$) in the 1000 Genomes Project. Statistical significance was assessed with the Wilcoxon rank-sum test.

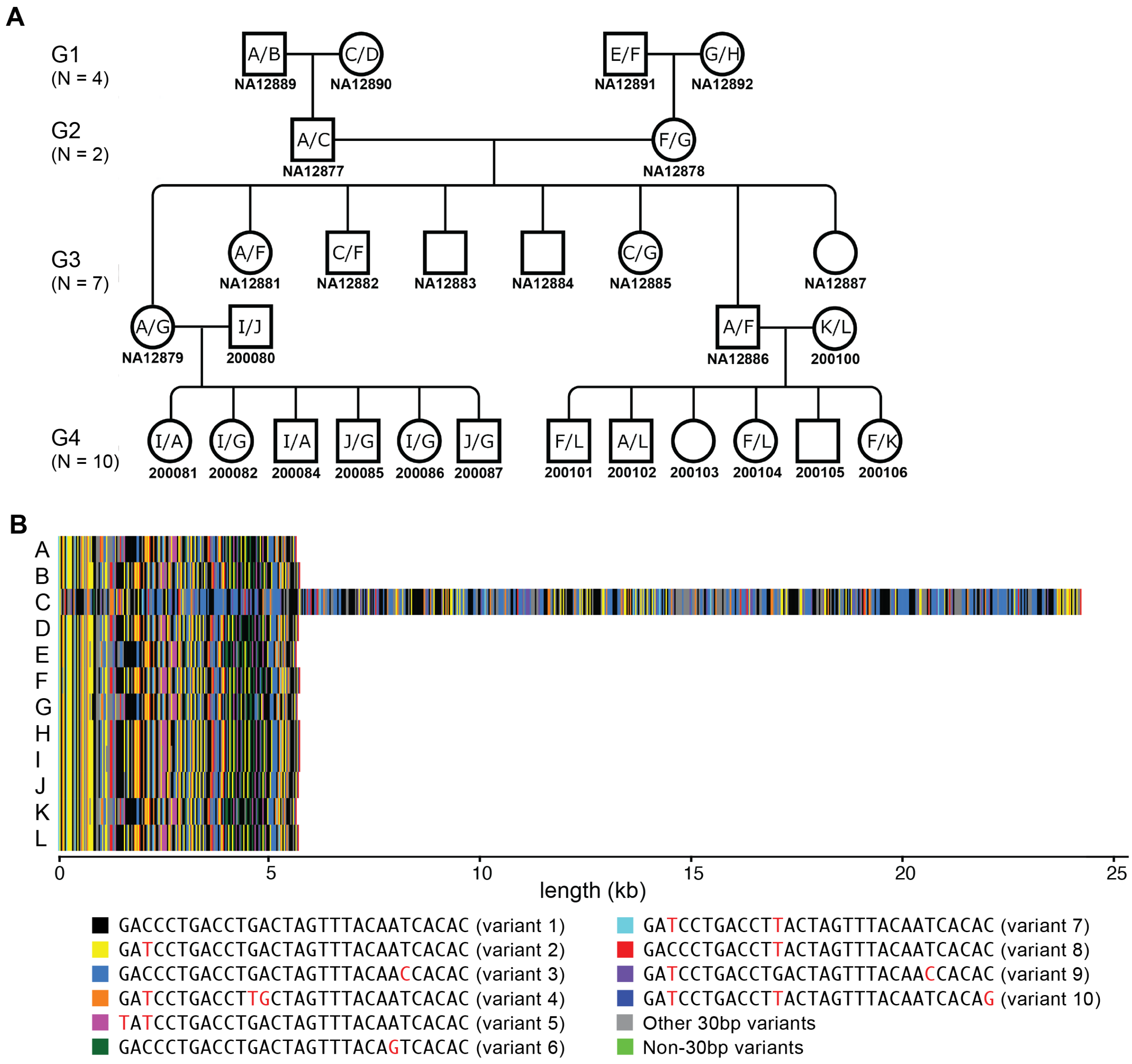


**Figure S5: No evidence for TRACT germline instability in the Platinum Pedigree.** (A) Alleles (labeled A-L) segregate as expected in a four-generation pedigree. No variation in TRACT length was detected. Data were not readily available for unlabeled individuals in the pedigree. (B) Visualization of all alleles found in the pedigree, with coloring of the ten most common variants.

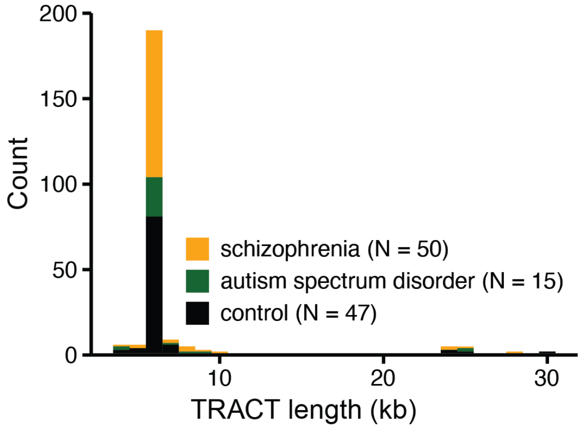


**Figure S6: Distribution of TRACT length in individuals assayed for somatic mosaicism.** Stacked distribution of TRACT allele lengths for 47 controls (black), 50 individuals with SCZ (orange), and 15 individuals with autism (dark green), as assessed by Southern blot.

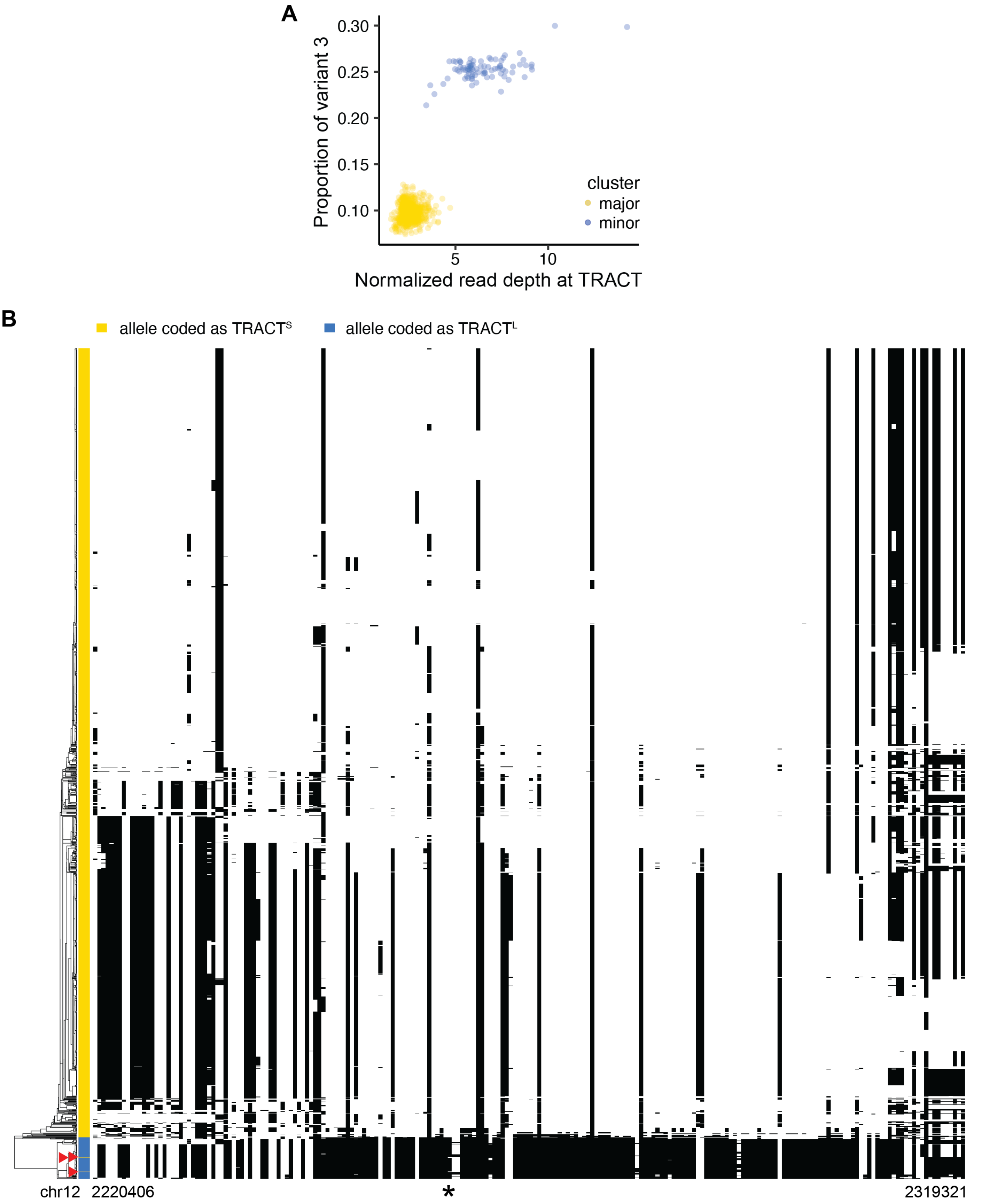


**Figure S7: TRACT$^L$ is found on one haplotype in GTEx.** (A) Individuals in GTEx (Lonsdale et al., 2013) with high variant 3 proportion are enriched for increased read depth ($p < 10^{-65}$, 2-sample Kolmogorov–Smirnov test). The x-axis is the number of reads that map to TRACT divided by the total number of reads in each sample $x10^6$. (B) TRACT$^S$/TRACT$^L$ was coded as a variant, and the 2 Mb interval surrounding TRACT was phased using Beagle 5.0 (Browning and Browning, 2007) (Materials and Methods). The phased alleles were visualized using Haplostrips (Marnetto and Huerta-Sánchez, 2017) for chr12: 2220406-2319321 (hg38). The genomic position of TRACT is indicated by an asterisk. Three red arrowheads indicate alleles from individuals that are likely homozygous for TRACT$^L$.

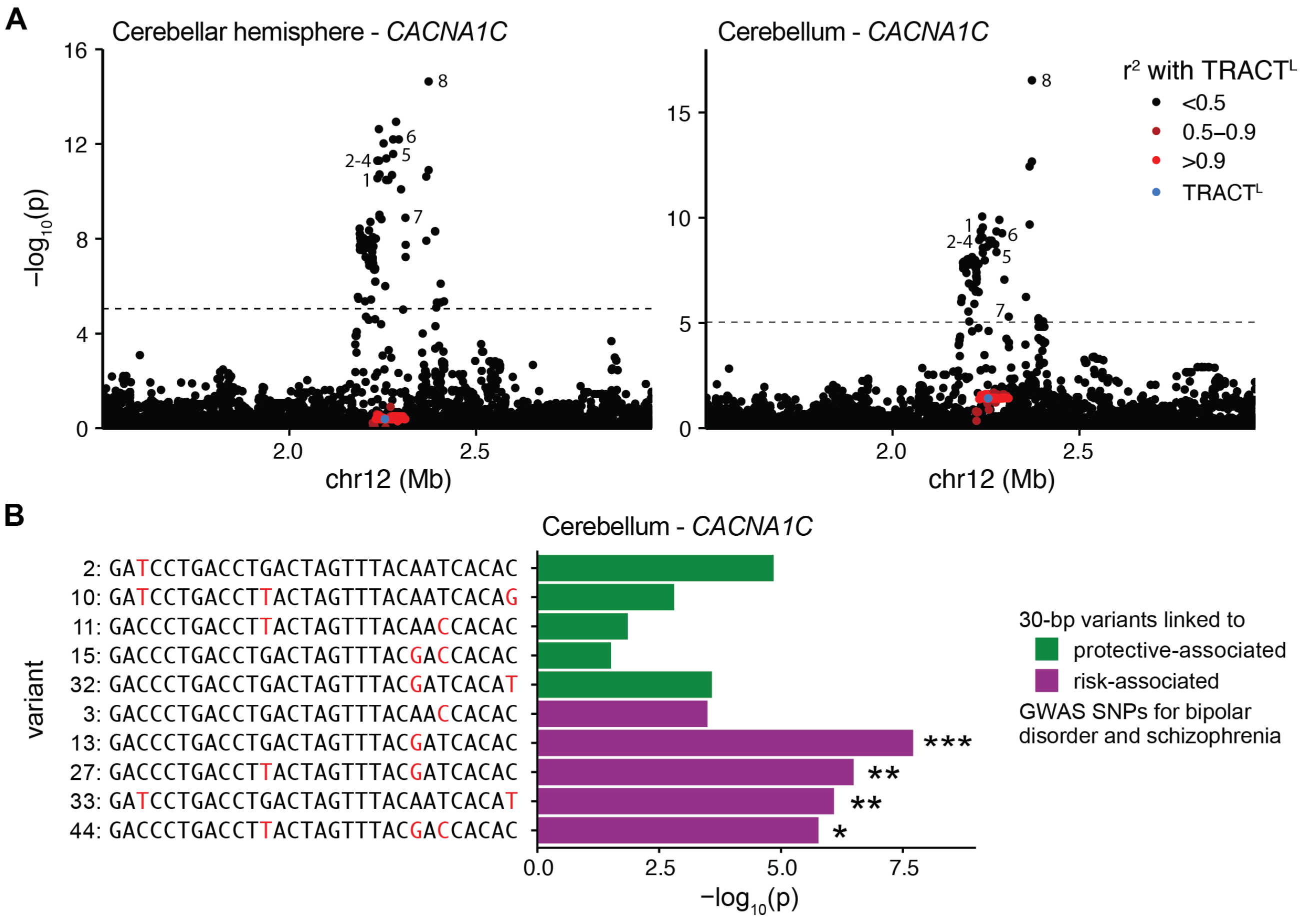


**Figure S8: Neuropsychiatric GWAS SNPs and specific 30-bp variants are eQTLs for *CACNA1C* in the cerebellum.** (A) The cerebellum was sampled twice in GTEx, labeled as either the cerebellar hemisphere (left) or the cerebellum (right). SNPs surrounding TRACT, but not TRACT$^L$ itself (blue point) or SNPs in linkage disequilibrium with TRACT$^L$ (red points), are eQTLs for *CACNA1C* expression in the cerebellar hemisphere (left) and cerebellum (right). SNPs labelled 1-7 are previously identified GWAS SNPs (Ferreira et al., 2008; Ripke et al., 2011; Sklar et al., 2011; Smoller et al., 2013; Ripke et al., 2013, 2014; Ruderfer et al., 2014; Song et al., 2018; Ikeda et al., 2019; Stahl et al., 2019; Lam et al., 2019). SNP 8 is an eQTL for *CACNA1C* in the cerebellum that is not in linkage disequilibrium with the GWAS locus. Fine mapping by GTEx suggests that the SNP 8 eQTL locus is distinct from the GWAS eQTL locus (Materials and Methods). 1: rs2007044, 2: rs1006737, 3: rs2159100, 4: rs4765905, 5: rs10744560, 6: rs1024582, 7: rs4765913, 8: rs886898. The dotted line indicates the significance threshold after Bonferroni correction for association with *CACNA1C* expression. (B) The proportion of particular 30-bp variants (13, 27, 33, and 44) are eQTLs for *CACNA1C* expression in the cerebellum. *: adjusted $p < 0.05$; **: adjusted $p < 0.01$; ***: adjusted $p < 0.001$.
